# Neural Networks as Entropic Systems: Applications in Digital Pathology

**DOI:** 10.64898/2026.01.30.702864

**Authors:** Alejandro Leyva, M. Khalid Khan Niazi

## Abstract

Deep learning systems in digital pathology are widely regarded as opaque, limiting clinical trust and interpretability. We present a framework for empirically characterizing training-time learning dynamics in neural networks by directly measuring activation structure, weight evolution, and spectral organization during optimization. Using TCGA-BRCA whole-slide images with replication-timing–derived methylation proxies as regression targets, we trained a Vision Transformer and tracked its intra-epoch behavior across 20 epochs.

We observed reproducible structural signatures during training. Correlated groups of neurons formed stable activation modules whose modularity increased as training progressed, accompanied by a reduction in representation entropy of up to 60%. Weight trajectories exhibited bounded diffusion with progressively reduced variance, consistent with a damped stochastic process, and converged toward a stable stationary regime in later epochs. In image space, model attention systematically shifted from collagen-rich stromal regions in early epochs to basophilic, proliferative nuclear regions in later epochs, aligning with known histologic correlates of replicative stress.

These findings demonstrate that neural networks develop predictable, quantifiable internal structure during training that can be directly visualized and measured. Framing learning dynamics in terms of entropy, modular organization, and stochastic stabilization provides a practical, mechanistic lens for interpreting how pathology AI models acquire biologically meaningful representations.

## 1 Introduction

Deep learning models are increasingly deployed in digital pathology, yet their internal learning processes remain difficult to interpret, limiting clinical trust and adoption [1]. While training objectives are formally defined by loss minimization, the mechanisms by which neural networks reorganize internal representations during optimization are not directly observable [2]. Neurons within a layer do not act independently; their activations and parameter updates are statistically coupled through shared gradients and correlated feature learning, leading to identity drift across training epochs and complicating neuron-level interpretation [3,4].

Existing interpretability approaches provide only partial insight into these dynamics [5]. Attention maps offer an output-level visualization of salient image regions but do not reveal how internal representations evolve during training [6]. Representation-similarity methods such as Canonical Correlation Analysis (CCA), Single Vector Canonical Correlation Analysis (SVCCA), and Projection-Weighted Canonical Correlation Analysis (PWCCA) quantify alignment between layers or checkpoints but operate primarily as snapshot comparisons, obscuring the temporal structure of learning [7]. As a result, current tools are limited in their ability to characterize how features are consolidated, reorganized, or deprioritized as optimization progresses.

Because neuron activations are correlated, they can be meaningfully grouped into modules, analogous to network-based approaches used in gene and protein interaction analysis [8,9]. Tracking groups of correlated units rather than individual neurons mitigates the effects of identity drift and enables the study of learning as a collective, structured process. Tissue structures are hierarchically layered, and are composed of patches or regions that can be classified using descriptors like “tumor”, “benign”, “necrotic” or otherwise. Within these patches, are cellular neighborhoods that define the behavior of the local regions within a patch, for example, malignant epithelial cells tend to cluster and agglomerate to create barriers against the local immune system. Within these cellular neighborhoods are the cells, which can be leukocytes, stroma, epithelial cells, among others. When using neural networks, lLayers closer to the input or output often correspond to visually interpretable tissue features such as neighborhoods or cell structures, whereas intermediate layers encode more abstract statistical representations that are not directly accessible through standard visualization techniques, for example, subtleties or patterns in staining or topology [10,11]. Changes in these intermediate representations across epochs reflect shifts in feature prioritization driven by optimization rather than input variation alone. As the model learns which features minimize loss, those features will be considered by the model more than other features.

In parameter space, stochastic gradient updates can be approximated as diffusion processes under small-step, fixed-batch regimes, motivating a stochastic description of weight evolution during training [12,13]. These dynamics imply that learning trajectories exhibit structured variability rather than arbitrary noise. We show that If such stochastic structure is reflected in activation space, then internal representations should display measurable changes in organization, entropy, and stability as training proceeds. Importantly, these changes are expected to be observable without requiring access to individual neuron identities.

Here, we introduce a means to track training-time learning dynamics by modeling activation layers as correlative networks and quantifying their modular organization, entropy, and stability across epochs. By jointly analyzing activation structure, weight diffusion, and image-space attention, we provide a dynamic view of how neural networks acquire and consolidate biologically relevant features during optimization. Applied to whole-slide pathology models, this approach may enablees clinicians and computational pathologists to assess whether models learn interpretable, tissue-grounded representations and to diagnose training behavior from a mechanistic, cause-and-effect perspective.

## 2 Materials and Methods

This section describes the experimental setup and diagnostic pipeline used to characterize intra-epoch learning dynamics visually. The motivation was to measure activation dynamics and the motion of layers across the three spaces of Vision Transformers: activation space, parameter space, and image kernel space. Formal derivations and theoretical results are provided in the Supplementary Theory. All definitions and acronyms are summarized in the Glossary.

Activation modules were extracted by first constructing correlation matrices between individual neurons and then agglomerating groups of four to five neurons into modules based on correlation strength, extending established methods in representation analysis such as CCA, SKCCA, and SVCCA [14]. Each activation matrix was defined by a sample count and a neuron count, and correlation-based grouping allowed the activation layer to be treated as a network, following practices established in community detection and network modularity literature [11–14]. This enabled the tracking of neuron clusters rather than individual neurons, mitigating the effects of identity drift that arise from correlated weight changes during training [15].

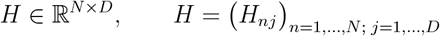

Z scoring of Pearson correlation and sequential top-k scoring was used to threshold edges and create modules, and a greedy community detection algorithm was then applied to form communities around the nodes, following standard modularity-based network approaches [16].

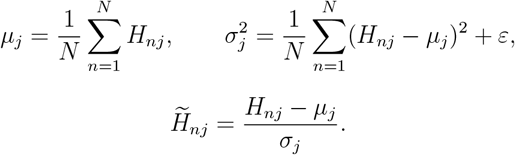

Pearson correlation coefficient is computed between neurons is computed as follows for j,k:

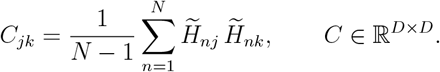

The edges, or correlative connections between nodes or modules, are Z-scored and thresholded using top-k:

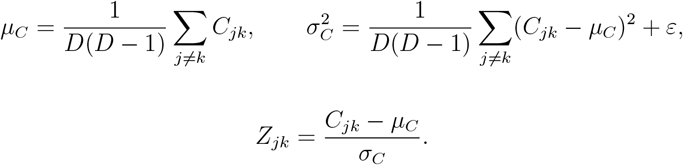

Sequential Top K is done as follows to measure the strength of each correlation, which is effective since most neurons have a large number of correlations.

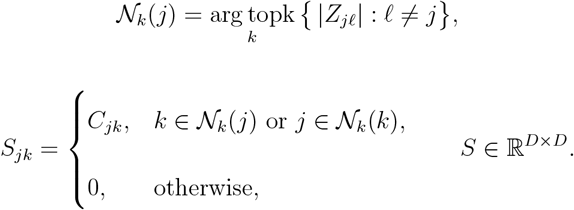

greedy communities is performed to agglomerate the modules using the top weighted edges between each neuron based on proximity:

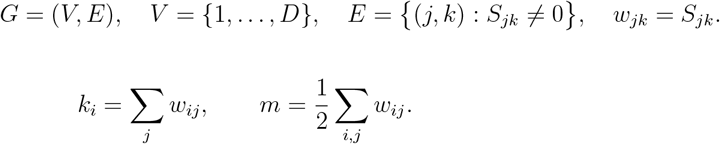

Greedy communities were evaluated using the Newman–Girvan modularity to measure the stability of the agglomerations [16,17].

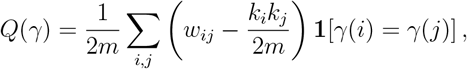

where *γ* : *V* → {1, …, *K*} is a community assignment. The greedy algorithm seeks 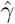 such that 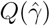 is locally maximal.

Maslov–Sneppen degree-preserving rewiring was used to form null communities as a control, ensuring that observed variations were due to genuine organization rather than chance [13].

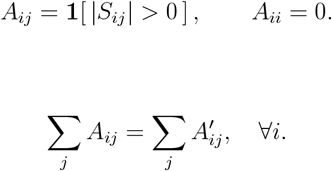

Rather than assuming a theoretical distribution, percentile bootstrap confidence intervals were used to resample edges and reconstruct the empirical distribution of edge weights and drift statistics [18].

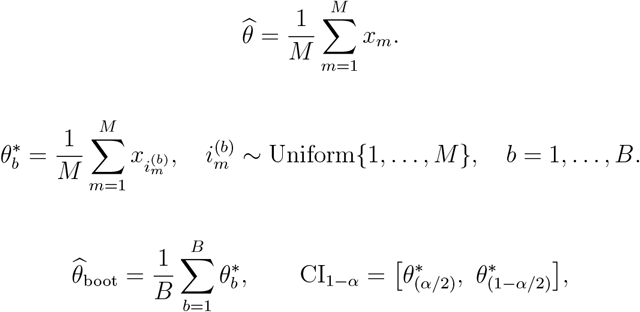

where 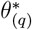 is the empirical *q*-th quantile of 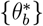.

The network constructions were compared against Laplacian-based graph metrics, Hungarian continuity matching, and representation-similarity methods including PKCCA and SVCCA to assess structural and representational stability [19].

Laplacian metrics were obtained from the weighted adjacency matrix and used to compute eigenvalues, from which a spectral entropy measure was derived using Shannon entropy to quantify the complexity of the connectivity spectrum [20].

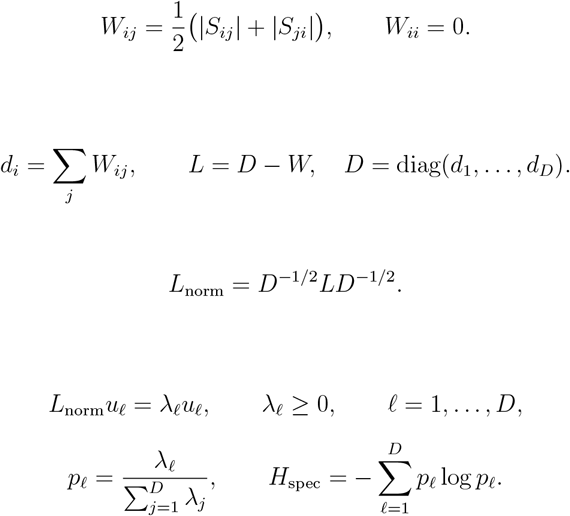

Hungarian continuity metrics were defined for each module and edge set to track module identity over epochs via optimal assignment, using the Hungarian method as the underlying matching algorithm [20].

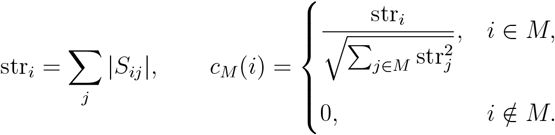

Centroids between each set of nodes are built along with the cosine similarity matrix.

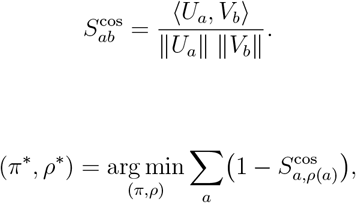

where *ρ* is a permutation of current modules. The jacquard stability is measured as follows

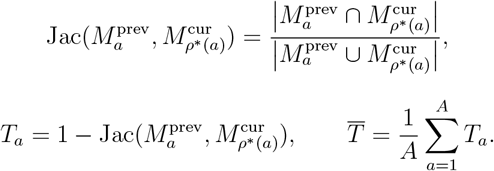

The drift of each module was done using CKA and PKCAA, where the identity drift and thus the stability of each module in the 3rd layer could be measured across epochs, since intra-epoch activations are difficult to track accurately due to inherent lack of bijectivity between image features[21]. The layer representations X and Y in the set R is characterized by the number of samples and the number of neurons that changes in each epoch.

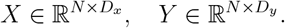

The columns are centered, and then correlationally computed to observe the change in correlations across epoch.

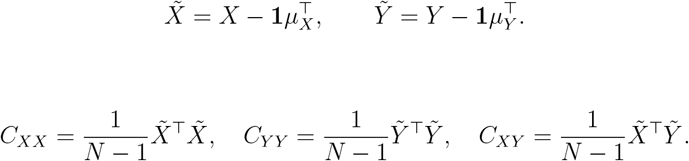

Where the cross covariance is computed, or the drift:

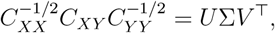

where the singular values on the diagonal of Σ are the canonical correlations *ρ*_1_, …, *ρ*_*r*_.

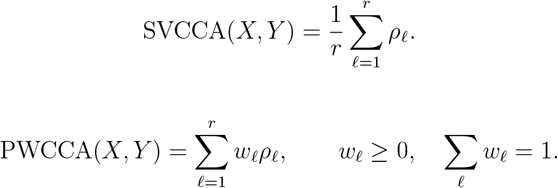

the top ranked patches and attention scores were recorded and an image was generated for each epoch. The graphs for the network were kept in 3d and 2d images for layer 3. modules were matched based on cosine similarity, and then tracked using Cosine similarities, hungarian tracking, null referencing, identity drift, modularity, and entropy across each epoch

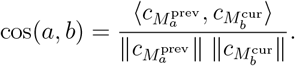

We begin with standard stochastic gradient descent assuming a constant learning rate aeta with fixed batch size S with no weight decay. Local basin and mini-batch sampling at small steps were used to simulate local drift within the epoch. The covarince matrix of each weight parameter theta of k is a summation, and the expectation of noise parameter epsilon is zero.

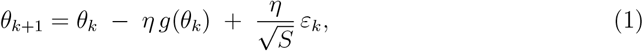

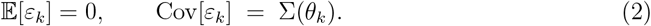

Using the definition of continuous time defined as the product of increasing integer set k and decreasing step, then under the central limit theorem, the change in weight parameters over time can be equated to change of theta. With Noise being a random variable, the expectation or mean randomness introduced can be summarized as a wiener process or W, whose covariance is the algebraic definition of matrix summation. using simplification and term rearrangement such that the square root of aeta over aeta is aeta, the terms can be arranged using the definition of the weiner process to simplify the equation as follows:

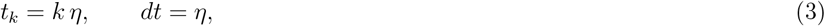

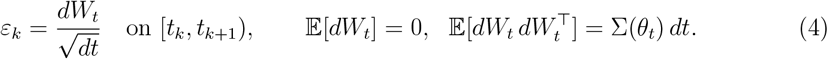

Now summarizing stochastic gradient descent using the wiener process, and using the defitition of covariance of the wiener process, the randomness of the process used to describe the change in weights over time can be simplified as follows:

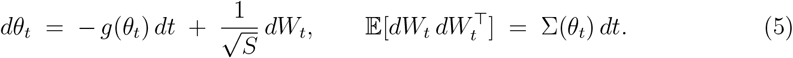

Using the definition of the weiner process with respect to brownian and diffusive processes, the equation can be simplified into a second order langevin equation with respect to flux B, diffusion D, and loss L.

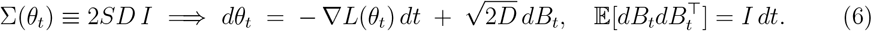

Diffusivity is defined using the following, where S is batch size and aeta is constant learning rate, assuming diffusivity is constant with respect to covariance.

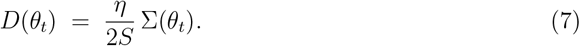

The langevin state equation using diffusion can then be simplified into the Fokker Planck equation, where the change in weights over time is dependent on the divergence of the product between the probability of weight at time t with loss g summated with the expanded diffusion term:

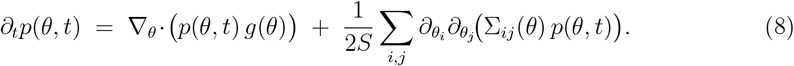

Simplifying terms for constant diffusion:

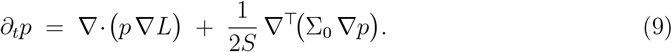

resulting in bayesian inferenced distribution of the likelihood of any weight being at a particular point.

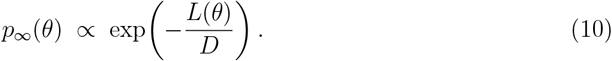

Assuming anisotropic diffusion, whereby the movement of each weight is not the same with respect to each other:

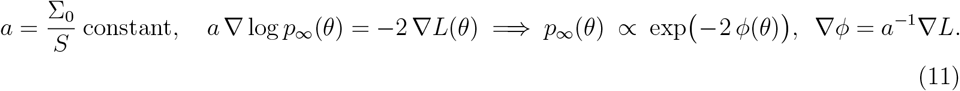

We can then describe this process over time as an ornstein uhlenbeck process to model the local effects of movement with respect to time

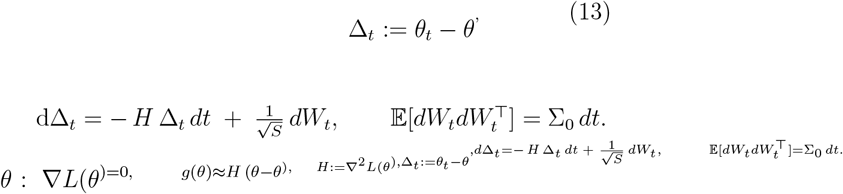

We can solve the lyupanov equation to solve for the constant covariance between two weights assuming diffusion is constant across learning rate:

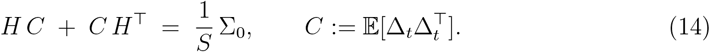

Solving for eigenvalues, and using the definition of C with respect to the change in weight positions over time, the change of one weight i can be modeled as a constant covariants dependent on the eigenvalue from the hessian equation

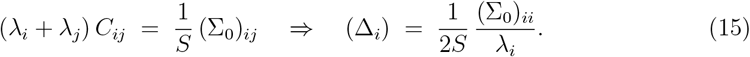

The equation for flux J is the difference between the probability of the change of weights over time with respect to loss and change in probability with respect to constant covariance alpha, where by the change in p can then be described as the change in flux using constant diffusion.

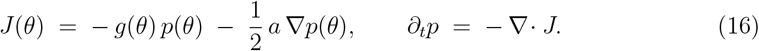

Entropy can be characterized using the weight parameter distribution using Gaussian distribution C, probability mu, batch noise epsilon, and the difference between dimension d and time step k.

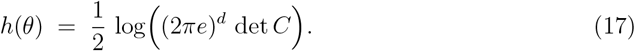

Within the network, the top eigenvalues or most prevalent weights can be determined as follows:

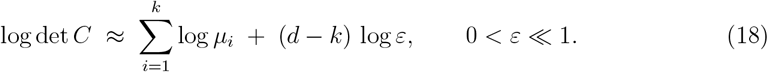

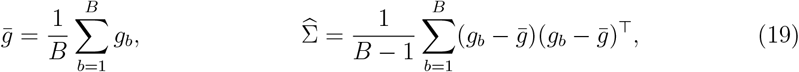

Effective diffusion within a single epoch can be modeled with temperature aeta and constant covariance.

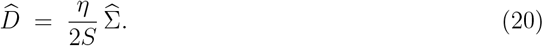

to measure the covariance across snapshots of time in the epoch, each weight can be measured across each batch size B and gradient g

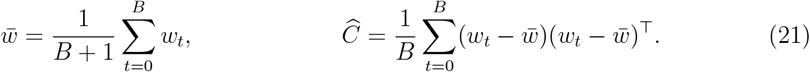

In order to measure across timesteps within the epoch, the covariance across each batch must be measured to ensure that the covariance remains staionary.

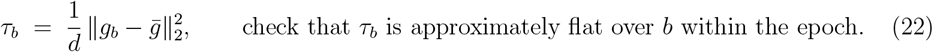

We begin with convolutions on each patch on the image H x W and patche embedding conv with weight W influenced by the Input channels (3 for RGB), output channels embedding dimension d (C, 256x 256 for ViT), Filter K for width and height whereby for an 8×8 patch for example would use KH x Kw.

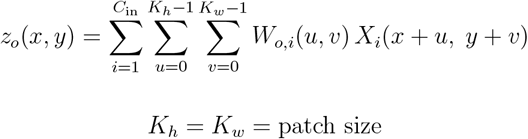

The weights for the 2D images are then spatially centered using the following equation, where the actual embedding weight is subtracted from the average learned weight across each row and column for mean brightness measurement to ensure that frequency is measured by shape rather than offset:

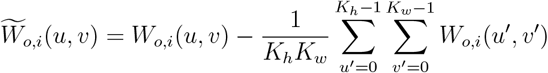

Using padding, the dimensionality can be set using the following equation, where p is the number of pixels padded around the image for focus:

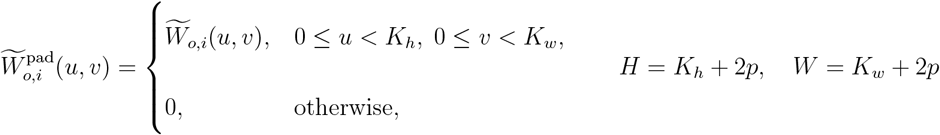

The filters are then transformed into the 2D fourier domain using double summation across all rows and columns and fourier transform across each centered weight omega. where k and l are the frequency indices measured across each row and each column.

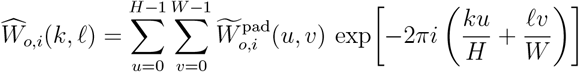

the power spectrum for a single filter is thus:

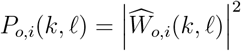

and the global power spectrum is the average power spectrum across both input and output channels:

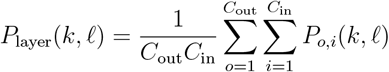

On the H x W frequency grid, the coordinates are centered on the lowest row and column, where the euclidean radius is described using polar coordinate scales, which can be normalized using the maximum radius against the number of bins.

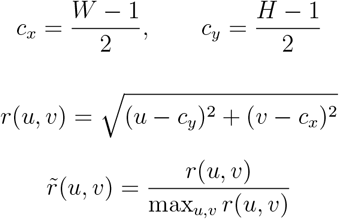

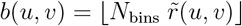

Total power is defined as a summation of power spectra and can become a probability distribution:

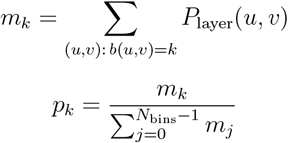

Using this probability distribution, shannon entropy can be measured throughout each training step t within the epoch, with a running mean to smooth the curve:

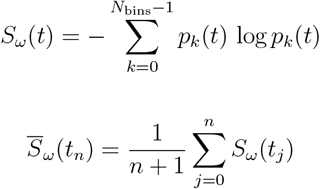

Also, the fourier transform across all patches was performed, the power spectra were taken, radially binned and normalized to form the probability distribution for the shannon entropy measurements, which was then averaged over batches. gradient trace was also applied with a running mean to ensure lower noise and observe boltzmann like conditions:

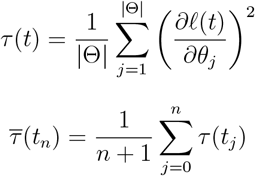

By this means, image spectral density can be tracked through intra-epoch with respect to changes in weights.

We trained a standard S-16 Vision Transformer (ViT) on whole-slide histopathology images from the TCGA-BRCA cohort to predict replication-timing–derived methylation proxies as continuous regression targets. Replication timing is defined as the replication rate of genetic domains within the interphase of cellular division. Domains that have a higher replication timing rate will typically be highly expressed, resulting in morphological changes reflecting oncogenesis or changes in metabolic rate. Predicting replication timing allows for gauging the activity of DNA damage and response genes and predicts the proliferative ability of the tumor. Replication timing is highly prognostic and infers several mutations within lung and breast cancer [22]. Importantly, methylation data were used only to derive slide-level labels reflecting replicative stress, as Repli-seq data is not available at a large scale. Thus, methylation-based proxies based on known correlations and mechanisms would reflect replication timing [23]. The; model inputs consisted exclusively of image patches extracted from whole-slide images, and were used to predict the replication timing proxy. Using a simple regression task based on biological labels, we isolate image features for prediction and exclude clinical covariates for isolation. This formulation enabled the study of learning dynamics in image-based models while grounding predictions in independent molecular measurements.

The dataset comprised 1,130 whole-slide images paired with corresponding methylation profiles, including idat and beta-value matrices used to compute replication-timing proxies. All preprocessing details to derive replication timing labels are available [24].The model was trained to predict replicative stress on a per-slide basis. Training was performed using Python, and experiments were executed on the Ohio Supercomputer Center (OSC) Ascend cluster for distributed computation. Code for the full pipeline is available upon request.

Since singular neurons cannot be tracked at an individual level, Neural dynamics can be measured using the property that neurons are not mutually independent in behavior. To characterize representational structure, activation modules were extracted by constructing correlation matrices between neurons within a given layer at each epoch. Using the networks, we can visualize the movement and dynamics of clusters of neurons in activation space, measure the distances traveled, measure the entropy across epochs, and observe feature prioritization without assuming bijection between a neuron and a particular feature. Groups of four to five highly correlated neurons were agglomerated into modules based on correlation strength, extending established representation-analysis techniques such as CCA, SKCCA, and SVCCA [25]. Each activation space matrix was defined by the number of samples and neurons, allowing activation layers to be treated as weighted networks.

This network-based formulation follows standard practices in community detection and modularity analysis [26] and enables tracking of groups of correlated units rather than individual neurons. While the linear correlation of these modules does not correspond to the similarity of the features they represent, it can measure how certain features covary with their change in weights due to backpropagation. It is important to note that neurons are filtered from the model if they have no correlation or edges with the global network. Aggregating neurons into modules mitigates identity drift arising from correlated weight updates during training and allows learning dynamics to be studied at a mesoscopic scale that is stable across epochs [27].

We also model learning dynamics using the same principles in parameter space, but rather than using network dynamics due to the size and scale of parameter space, we model the change in the distribution of weights within parameter space. Within an epoch, stochastic gradient updates with small step sizes and fixed batch structure can be approximated as diffusion processes, motivating a quasi-Boltzmann description of weight evolution [28]. Weight trajectories were therefore analyzed in parameter space using principal components to characterize diffusion, variance damping, and stabilization over training. This in turn provides a visual and interpretable understanding of learning dynamics, as changes in the weight distribution can be modeled entropically.

Because patch embedding operators in Vision Transformers are linear projections of image patches parameterized by model weights, changes in weight space induce corresponding changes in how local image structure is encoded prior to attention-based integration. While downstream transformer blocks introduce nonlinearity through attention and MLP layers, the patch embedding stage itself remains a linear operator, making its spectral properties directly interpretable. Accordingly, we model the distribution of spectral power in the patch embedding weight space across training epochs, treating embeddings as parameterized linear mappings rather than standalone feature detectors. Learned embedding kernels were analyzed in the frequency domain using classical Fourier and signal-processing formulations, from which spectral power distributions were derived [29]. Power spectra were computed from filter weights and summarized as probability distributions over frequency components. Shannon entropy was then used to quantify spectral organization and variability over time [30]. For the full derivation of the power spectra distribution, please see the supplemental section and the GitHub.

This joint analysis independently characterizes parameter-space diffusion, activation-space modular organization, and image-space spectral structure, offering a unified descriptive framework for visualizing and measuring learning dynamics across multiple representational spaces during training.

## 3 Results

The network structures shown in PCA space in Figure 1s demonstrated high modularity and entropy over time, with the 3D networks alternating structure across epochs. Each frame corresponds to one epoch, and the transition states intra-epoch are not modeled. Interestingly, similar network geometries recur toward later epochs, suggesting partial structural convergence rather than purely stochastic reorganization. For example, In epochs 11-12, the jaw like structure represents a stark division between neural modules, that recurs in epoch 17-18. For context, these behaviors are the result of the fact that different weights do not have the same change across each iteration of backpropagation. As a result, Modules with smaller weight changes agglomerate and so forth. These snapshots show the structure of the network after equilibration within each epoch. Network structure is most diffuse in epochs 1–3, whereas epochs 17–20 show higher modular stability and more coherent community organization.

**Figure 1.**
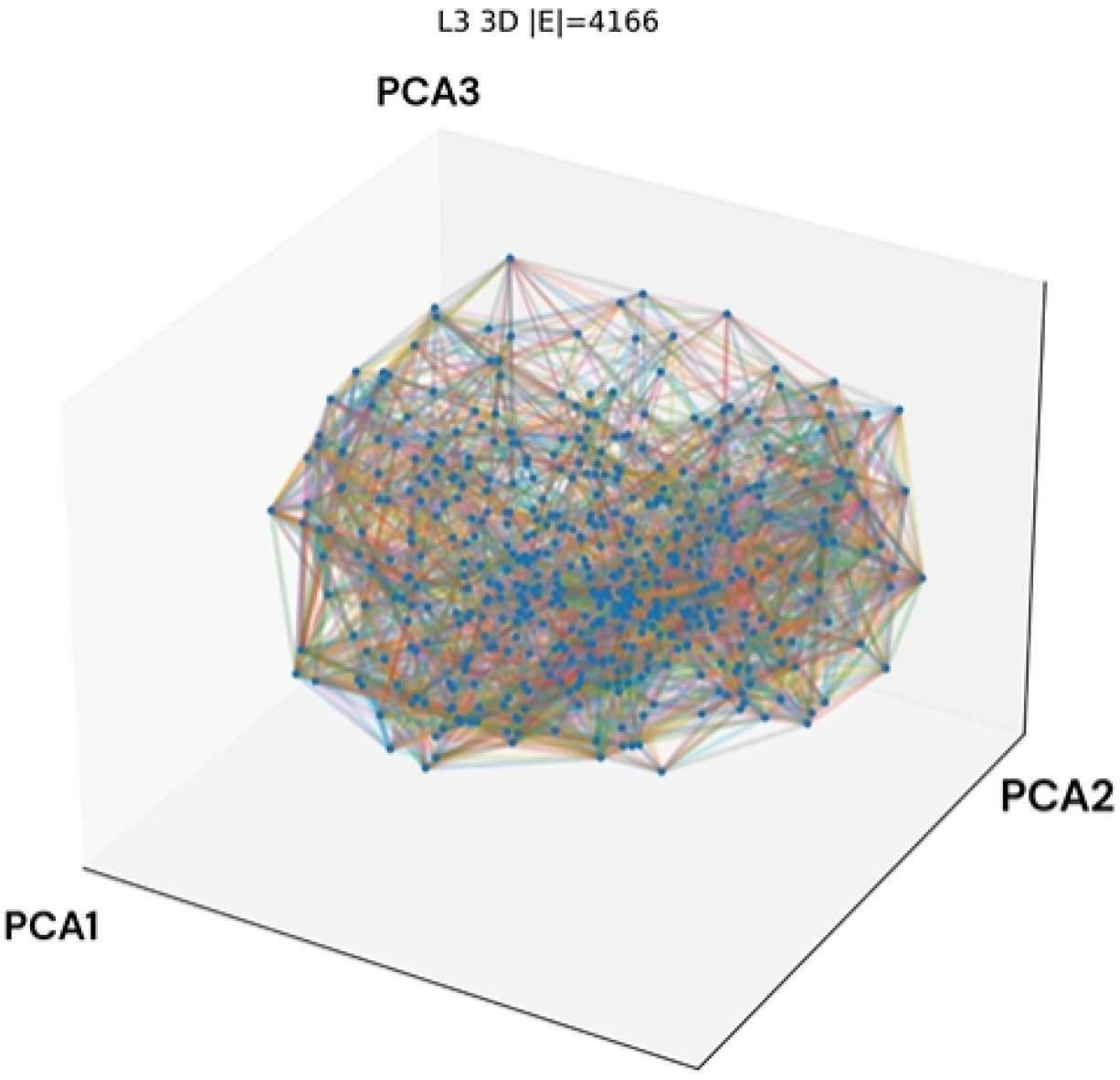
Layer L3 3D structure animation. PCA is ordered from highest to lowest.

Across the attention maps for the top modules shown in Figure 2, several basophilic glands recur, and in epoch 1, the patch that was ranked the lowest became the highest-ranked patch in epoch 20. Generally, the top attention matches are stable, with a few rank switches concentrated in epochs 10 and 11. Figure 1 demonstrates a high level of identity drift across epochs, though modularity and the number of modules remain relatively stable.

**Figure 2.**
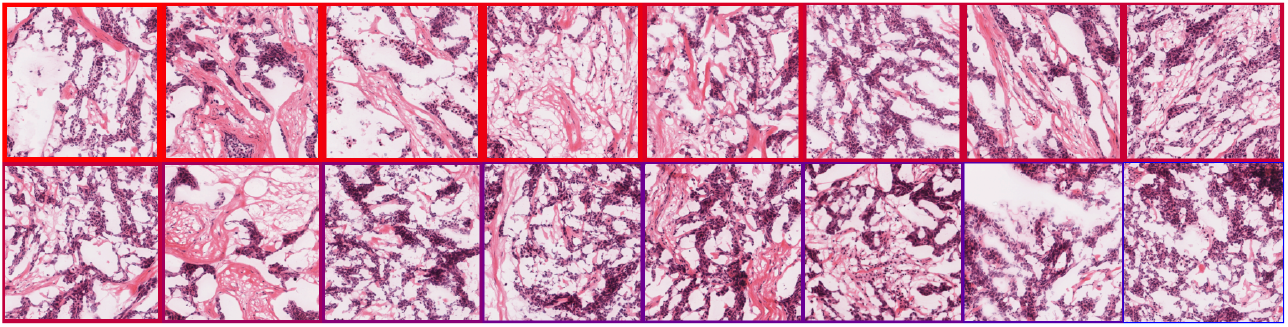
Layer L3 top-module animation (click to play).

Figure 2 shows changes in the top-ranked patches across epochs by ranking the attention weights. Notably, the lowest-ranked patch in epoch 1 became the most prioritized patch in epoch 20, while several patches either move two to three positions between epochs or matriculate out of the ranking entirely. Biologically, these epoch-wise changes correspond to a shift from prioritization of stroma-rich collagen regions in early epochs to basophilic, high-density nuclear regions in later epochs as sources of replicative stress. Basophilic nuclei frequently exhibit high replicative stress due to mesenchymal plasticity and transfer of oncogenes into early-replicating regions, resulting in higher and less regulated expression of factors such as WNT or MYC. While the specific image features detected by the model are not directly observable at the neuron level, the learning dynamics and their alignment with known biological mechanisms provide a means to ground model training.

Figure 3 shows the weight-parameter distribution trajectory across principal components using gradient vectors by flattening and normalizingin gradient space, where the distribution oscillates within a bounded range. Gradient tracing also shows that variability diminishes over time as a damped system, indicating that modular organization improves as the epoch concludes. Loss is variable per batch early in the epoch and is minimized toward the latter end. While diffusion of weight parameters may be damped due to the averaged effects of neural drift, this behavior provides a means to understand parameter space as a semi-open entropic system. Here, t refers to the intra-epoch timestep, computed based on identity drift over time.

**Figure 3.**
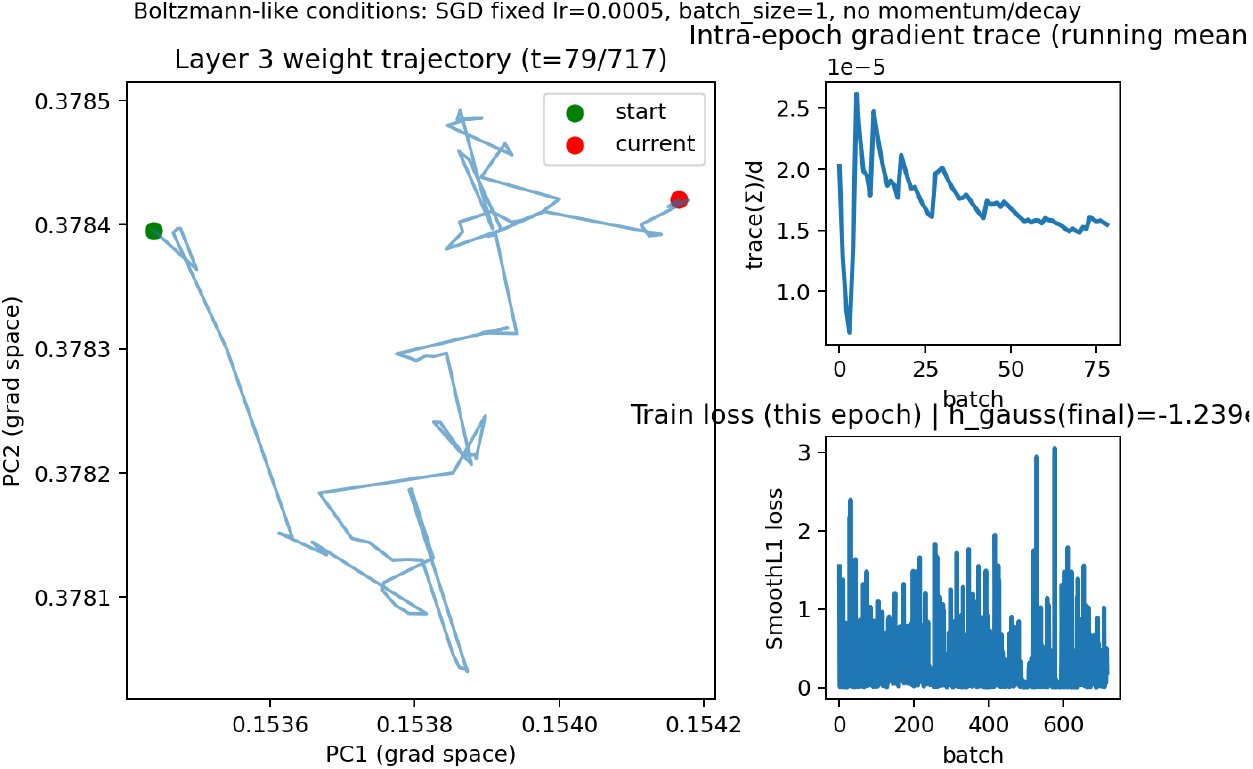
Layer L3 top-module animation (click to play).

Figure 4 visualizes effects of weight-space dynamics in the frequency and image-processing domains, where spectral entropy is measured from the power distribution of Fourier-derived weights. Because ViT patch embeddings are implemented as strided 2D convolutions, their weights define an explicit spatial filter bank; spectral entropy was therefore computed from the 2D Fourier power spectra of the patch-embedding filters. Global spectral entropy declines as the epoch concludes, although the effect size is small (∼ 10^−6^), while training loss stabilizes near a minimum toward the middle-to-late portion of the epoch. Gradient trace indicates that Boltzmann-like conditions remain fairly stable, and this stability is consistent with the minimal but systematic changes observed in spectral entropy. The filter spectrum remains consistent with low frequencies centered near zero and expanding outward. Reference spectral input entropy should remain constant, indicating that entropy changes observed in the image-processing domain originate from the model rather than the input distribution. Individual filter-output spectral entropy also demonstrates a consistent decline across epochs.

**Figure 4.**
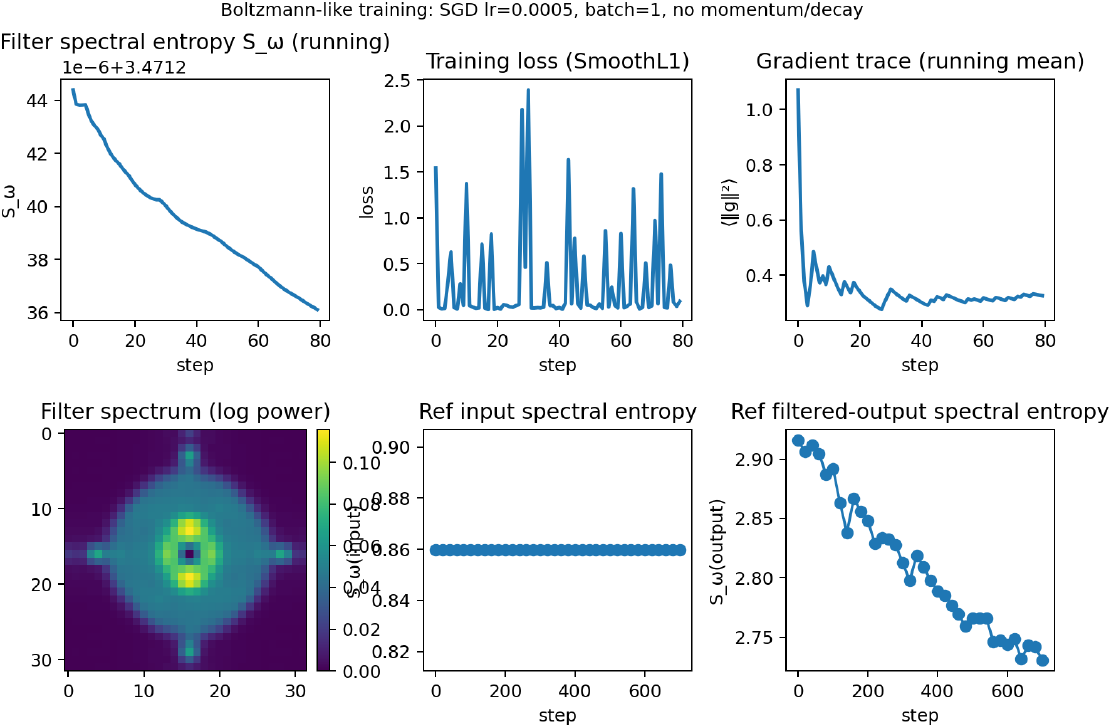
Layer L3 top-module animation (click to play).

Figure 5 shows representation dynamics for layers 1, 3, and 5 across 20 epochs. Panel (a) shows a subtle decrease in training loss and correspondence between training and validation loss, supporting the observation that entropy decreases as loss is minimized. Panel (b) shows that modularity is not constant across epochs and is heavily variable due to identity drift of activations; this is observable through shifts in top-ranked modules and changes in network structure, despite constant node counts and consistent significance relative to Maslov–Sneppen constructions. Panel (c) indicates a spike that dramatically decreases representational similarity in epoch 11. Structurally, epoch 11 is uniquely chaotic, with large node agglomerations and sparse bridging nodes forming a jaw-like structure. Otherwise, most epochs and adjacent layers show decent consistency. Epoch 11 also coincides with an increase in validation loss, while training loss marginally increases. Representation similarity is additionally supported by the module stability trend shown. Panel (d) shows a higher fraction of modules participating in the network and shifting spatially, indicating that a large proportion of nodes contribute to network reconfiguration. Weight turnover remains consistently high until epoch 16, where many nodes occupy positions similar to the previous epoch, serving as a transition state between epochs 15 and 17. Epochs 15–17 also show stagnation of validation loss. While similarity between adjacent layers remains high and modularity remains unstable due to high weight turnover, representation entropy decreases dramatically across epochs, suggesting that later epochs consolidate information learned earlier in training.

**Figure 5.**
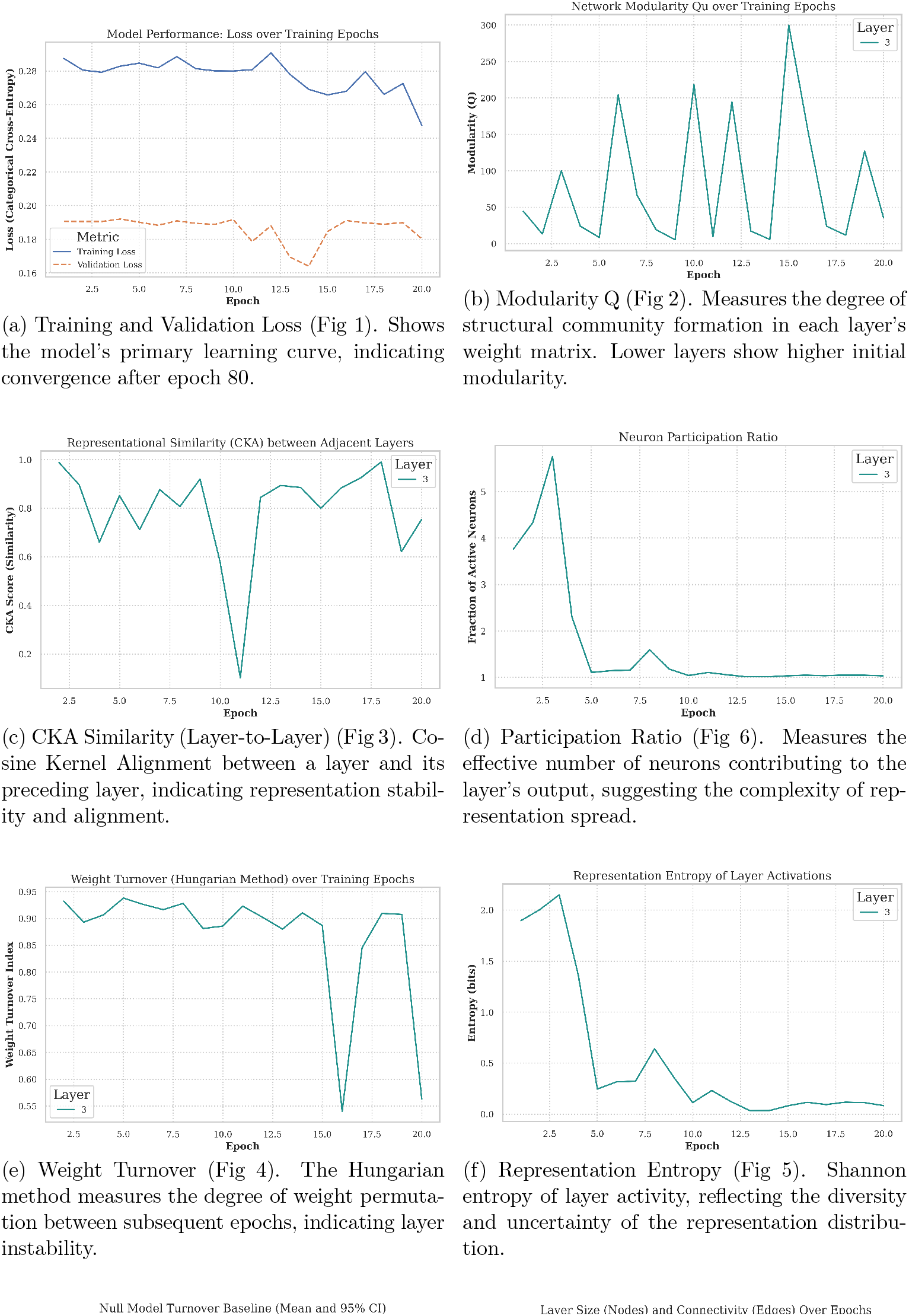
Core Performance and Representational Dynamics Over Training. This figure tracks basic learning metrics, layer stability, and representation quality across epochs.

Figure 6a shows a steady decline in edge entropy, or the likelihood changes in the number edges over time, and, concurrently, a steady decline in the maximum eigenvalue derive from the Laplace–Beltrami transform, likely because epochs stabilize as loss decreases, even though weight turnover remains consistently high. Using SVCCA and PWCCA, the same trend occurs, where epoch 11 shows very low representational similarity, indicating that these comparisons support the same conclusions. The overall number of nodes per layer and the number of edges remain relatively stable across epochs. The dotted line shows the number of edges, which was scored and filtered appropriately to model the top nodes. Based on panel (e), lower weight velocity (identity drift) corresponds to higher modularity, and panel (f) shows that increasing weight velocity does not necessarily imply higher entropy or higher turnover. In this system, which is not strictly modeled under Boltzmann equilibrium conditions, it remains plausible that some weights change substantially more than others.

**Figure 6.**
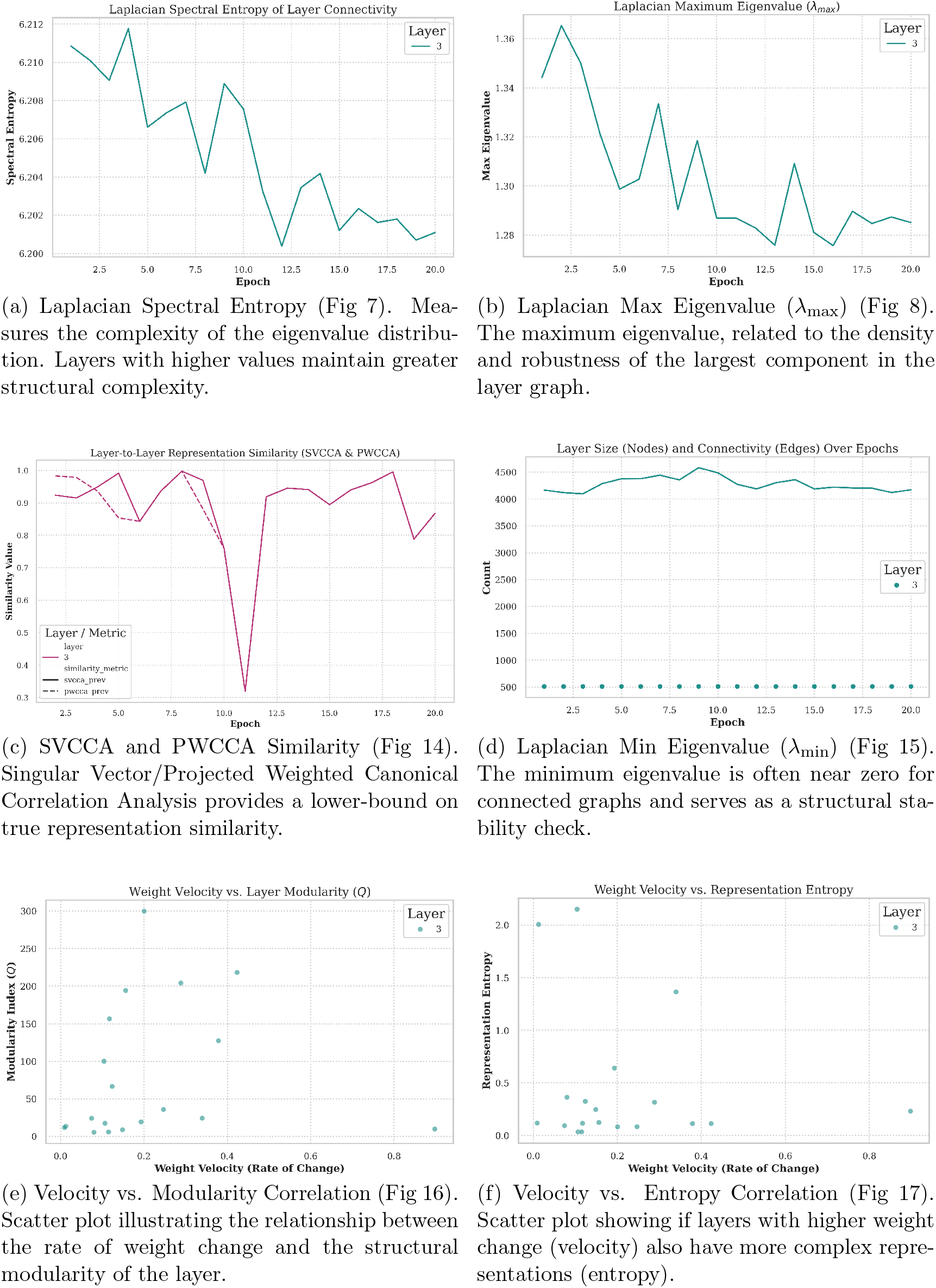
Advanced Spectral Metrics and Correlation Analysis. These figures explore the deep structural properties of the layer weights and the interaction between different dynamic metrics.

It is clear that intra-epoch learning dynamics and diagnostics can be visualized to understand the process by which features are valued and consolidated during training.

## 4 Discussion

Neural network training dynamics can be meaningfully approximated using stochastic frameworks without assuming that models operate as closed or equilibrated physical systems. While strict equilibrium is not attainable during optimization, equilibrium-inspired concepts provide a useful descriptive language for understanding how networks progressively minimize loss and stabilize internal representations. In this study, we observed systematic reductions in representation entropy and weight variability both within epochs and across training, indicating that learning proceeds through structured consolidation rather than uncoordinated drift.

These dynamics offer a mechanistic perspective on common model behaviors. Changes in entropy, modular organization, and weight diffusion change reflect how features are prioritized, refined, or deprioritized through weight changes during training to minimize loss and may help explain how these spaces behave under phenomenaphenomena such as over-fitting, undertraining, and sensitivity to hyperparameters. Importantly, these effects emerge at the level of correlated neuron groups rather than individual units, reinforcing the need for mesoscopic analyses that are robust to weight changes. identity drift.

Figures 1–6 show that neural network training involves coordinated but non-monotonic reorganization across parameter, activation, and image-processing domains, even as loss decreases smoothly. Early epochs exhibit diffuse network structure, high representational entropy, and elevated weight turnover. As training progresses, internal metrics evolve unevenly, revealing transient restructuring events that are not captured by loss alone.

Activation-space analyses demonstrate that neurons form correlated modules whose membership and spatial organization shift across epochs. Despite substantial identity drift, the number of modules and nodes remains stable, indicating that learning proceeds through re-configuration of connectivity rather than changes in network size. Representational similarity measures identify a pronounced disruption around epoch 11, where adjacent layers temporarily lose alignment and modular structure becomes irregular, coinciding with increased weight turnover and a brief increase in validation loss.

Parameter-space analyses show that weights continue to move throughout training but within a bounded region. Gradient-space projections reveal anisotropic diffusion with progressively reduced variance, consistent with damped stochastic dynamics rather than static convergence. Importantly, parameter velocity, modularity, and entropy are only partially coupled: high weight velocity can occur in both high- highand low-entropy regimes, and increased modularity does not uniquely determine update magnitude.

Image-space analyses indicate that these internal dynamics correspond to systematic changes in image processing behavior. Spectral entropy of patch-embedding filters decreases over training while reference input entropy remains constant, indicating model-driven reorganization. Attention patterns shift from stromal regions in early epochs toward basophilic, nuclear-dense regions in later epochs, consistent with known histologic correlates of replicative stress.

In digital pathology, interpretability is essential for clinical trust and deployment. Existing tools primarily emphasize output-level explanations, whereas the approach presented here focuses on how representations form and stabilize during training. By visualizing learning as a dynamic process, clinicians and computational pathologists can assess whether models converge toward biologically grounded features and can better understand the limitations and failure modes of AI-driven predictions. This training-time perspective complements existing interpretability methods and provides additional context for evaluating model reliability in healthcare settings.

## 5 Limitations

Neuron tracking in this framework is inherently approximate rather than exact. While weight distributions and activation correlations can be reliably modeled at the group level, individual neurons cannot be uniquely identified or followed across epochs there is no bijection between a neuron and a feature.due to identity drift and the high dimensionality of parameter space. Correlation-based thresholding enables the tracking of dominant activation modules, but weaker connections necessarily lose resolution, limiting fine-grained interpretability.

Additionally, the scale of Vision Transformer architectures constrains direct visualization of full activation layers. Although parameter-space dynamics are more tractable in smaller models, extending exact neuron-level tracking to large-scale architectures remains an open challenge. The stochastic approximations employed here rely on fixed learning rates and batch structures within epochs; deviations from these assumptions may alter the observed dynamics. Future work is required to assess the generality of these diagnostics across architectures, optimization regimes, and clinical tasks.

## 6 Conclusion

This study demonstrates that neural network learning dynamics can be empirically characterized using statistical and network-based diagnostics that operate across parameter, activation, and image representations. By modeling activation layers as correlative networks and analyzing entropy, modularity, and diffusion during training, we show that deep models develop structured, predictable internal organization over time.

Applied to digital pathology, this framework provides a mechanistic view of how biologically meaningful features emerge during optimization, linking training dynamics to observable histologic patterns. These methods offer a foundation for clinician-facing diagnostics that move beyond static explanations and toward dynamic assessments of model behavior. More broadly, the approach establishes a practical pathway for interpreting deep learning systems through their learning processes, supporting the development of transparent and trustworthy AI in biomedical applications.

## 7 Ethics Statement

All datasets are public and anonymized.

## 8 Author Contributions

All work was performed by the first author, with supervision by the second.

## 10 Data Availability

TCGA BRCA is a publicly available Breast cancer dataset. All code is available in https://github.com/Alejandro21236/NetworkMachines-, and replication timing is available here https://github.com/Alejandro21236/ReplicationTiming

## Supplementary: Proofs and Formal Statements

### 12 Use Of Generative Artificial Intelligence

The author used ChatGPT for grammar, punctuation, and formatting.

=

#### Setup

Let *L*(*θ*) be the empirical risk for parameters *θ* ∈ ℝ^*d*^, and let *g*(*θ*) = ∇_*θ*_*L*(*θ*). Per-batch gradients are *g*_*b*_(*θ*) with 𝔼[*g*_*b*_(*θ*)] = *g*(*θ*) and Σ(*θ*) = Cov[*g*_*b*_(*θ*)]. Activations at layer *ℓ* for input *x* are 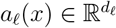 with forward map *a*_*ℓ*_ = *ϕ*_*ℓ*_(*W*_*ℓ*_*a*_*ℓ*−1_), where *W*_*ℓ*_ are layer weights and *ϕ*_*ℓ*_ is (piecewise) smooth.

##### Non-degenerate model

We assume (i) the network contains at least two coupled parameters and two coupled units per layer, (ii) the loss and architecture are not separable across coordinates, so the Hessian 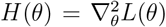 is not diagonal and some backprop Jacobians have nonzero off-diagonal entries.

###### Proposition 1 (Weights are not mutually independent)

*Unless the model and loss are fully separable across coordinates, weight updates are statistically dependent within an epoch*.

Consider one SGD step (fixed learning rate *η* and batch size *S*):

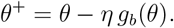

Let Δ*θ* = *θ*^+^ − *θ* = −*η g*_*b*_(*θ*). The covariance of updates is

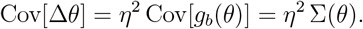

If weights were mutually independent in their updates, Cov[Δ*θ*] would be diagonal. Under the non-degeneracy assumption, backprop couples coordinates, so Σ(*θ*) has nonzero off-diagonals. Hence Cov[Δ*θ*] has nonzero off-diagonals and coordinates are dependent. Equivalently, at the level of expected drift, *H*(*θ*) having off-diagonals implies coupled deterministic motion, and at the noise level, off-diagonal Σ implies coupled stochastic motion. Therefore weights are not mutually independent.

###### Proposition 2 (Activations are not mutually independent and are rankable by correlation/connectivity)

*For a fixed distribution of inputs, within a given layer ℓ, the activation vector a*_*ℓ*_ *has a non-diagonal covariance matrix generically, and units can be ranked by correlation strength and graph connectivity*.

Write *z*_*ℓ*_ = *W*_*ℓ*_*a*_*ℓ*−1_ and *a*_*ℓ*_ = *ϕ*_*ℓ*_(*z*_*ℓ*_). If *ϕ*_*ℓ*_ is linear in a neighborhood and *a*_*ℓ*−1_ has covariance 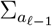, then

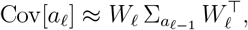

which is non-diagonal unless *W*_*ℓ*_ is a signed permutation and 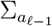 is diagonal in the same basis. With nonlinear *ϕ*_*ℓ*_, the delta method yields 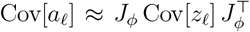 with *J*_*ϕ*_ diagonal only if all units are at identical slopes; generically Cov[*a*_*ℓ*_] stays non-diagonal. Hence activations are not mutually independent.

Define the empirical correlation matrix *R*_*ℓ*_ by normalizing Cov[*a*_*ℓ*_] to unit variances. Build an undirected weighted graph *G*_*ℓ*_ = (*V*_*ℓ*_, *E*_*ℓ*_) with vertices the units and edge weights *w*_*ij*_ = |*R*_*ℓ,ij*_| (or thresholded). Standard centralities (degree/strength *s*_*i*_ = ∑ _*j*_ *w*_*ij*_, eigenvector centrality, betweenness) induce a total preorder on *V*_*ℓ*_. Therefore units are rankable by correlation strength and by graph connectivity.

###### Theorem 3 (Boltzmann stationary law holds under constant diffusion, conservative drift, and small-step regime)

*Consider the continuous-time limit of SGD in parameter space*

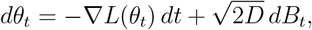

*where D is a constant scalar (equivalently* Σ(*θ*) ≡ 2*SD I) and B*_*t*_ *is standard Brownian motion. If the drift is conservative (*−∇*L), then any stationary density that satisfies detailed balance is*

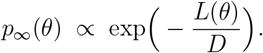

The Fokker–Planck equation is

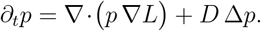

At stationarity and under detailed balance (zero probability current), 0 = ∇· *p*_∞_ ∇*L* + *D* ∇*p*_∞_, which for smooth, positive *p*_∞_ integrates to *D* ∇ log *p*_∞_ = −∇*L*. Hence log *p*_∞_ = −*L/D* + const, i.e., *p*_∞_ ∝ *e*^−*L/D*^.

*Remarks*. If diffusion is constant but anisotropic (*dθ*_*t*_ = −∇*L dt* + *GdB*_*t*_ with *GG*^⊤^ = 2*A* constant), then *p*_∞_(*θ*) ∝ *e*^−2*ϕ*(*θ*)^ with ∇*ϕ* = *A*^−1^∇*L*; if *A* depends on *θ* or if momentum/normalization layers induce non-conservative drift, detailed balance fails and the simple Boltzmann form need not hold. The diffusion approximation itself requires small learning rate, fixed batch size within the epoch, and i.i.d. sampling.

###### Theorem 4 (Entropy decreases as modularity/within-block correlation increases, under fixed marginals)

*Let X* ∈ ℝ^*d*^ *be zero-mean Gaussian with covariance C and fixed diagonal* diag(*C*) = *σ*^2^ *(fixed marginal variances). The differential entropy is*

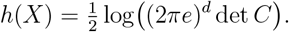

*Then h*(*X*) *is maximized when coordinates are independent (C diagonal). In particular, for a fixed community partition, increasing within-community correlations (i*.*e*., *stronger modularity) strictly reduces* det *C and thus reduces h*(*X*).

By Hadamard’s inequality, for any positive definite *C*,

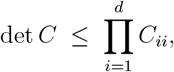

with equality iff *C* is diagonal (i.e., coordinates are uncorrelated). Fixing the diagonal entries 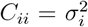, the product on the right is constant, so det *C* is maximized at independence and strictly decreases as off-diagonal magnitudes grow. Stronger modularity means larger within-block correlations (larger |*C*_*ij*_| for *i, j* in the same community) while keeping the diagonal fixed; hence det *C* strictly decreases and *h*(*X*) strictly decreases.

#### Corollary (Modularity vs. entropy, graph view)

Let *R* be the correlation matrix of *X* (unit diagonal). The Newman–Girvan modularity *Q* for a fixed partition increases as within-block |*R*_*ij*_| increase and cross-block entries decrease. Since 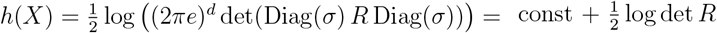, and det *R* decreases with larger within-block correlations, we obtain *Q* ↑ ⇒ det *R* ↓ ⇒ *h* ↓ .

#### Optional local OU characterization (for diagnostics)

Near a minimizer *θ* with Hessian *H* ≻ 0 and constant diffusion (within an epoch),

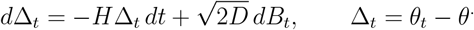

The stationary covariance *C* solves *HC* + *CH*^⊤^ = 2*D I*, giving Var(⟨*v*_*i*_, Δ_*t*_⟩) = *D/λ*_*i*_ along Hessian eigenvector *v*_*i*_ with eigenvalue *λ*_*i*_ > 0. Thus flatter directions (smaller *λ*_*i*_) exhibit larger equilibrium variance; adding correlation (modularity) lowers det *C* under fixed marginals, consistent with Theorem 4.

#### Summary of conditions for the Boltzmann claim

- Fixed learning rate within the epoch (small step), fixed batch size; i.i.d. batches.
- Conservative drift −∇*L* (no momentum/preconditioning in the claim, or incorporate them explicitly).
- Constant (locally) diffusion tensor within the epoch; isotropic for the simple *e*^−*L/D*^
- form.
- Local basin (linearization valid) for OU/Lyapunov diagnostics.

#### Standing assumptions (intra-epoch, late training)

Small step size; fixed batch size; plain SGD; i.i.d. batches. For local results, linearize around a basin point *θ* with *H* := ∇^2^*L*(*θ*^)≻0^. Gradient-noise covariance is locally constant: Σ(*θ*) ≈ Σ_0_.

[Non-diagonal Hessian ⇒ coupled deterministic drift] Let *L* : ℝ^*d*^ → ℝ be twice continuously differentiable. Consider the gradient flow 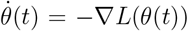 and its linearization near *θ* with *H* = ∇^2^*L*(*θ*^)^. If *H* has any off-diagonal entry *H*_*ij*_≠ 0 for some *i*≠ *j*, then the *i*-th coordinate of the drift depends on (at least) the *j*-th coordinate; in particular, the deterministic dynamics are not coordinate-wise independent.

Linearizing at *θ* with Δ(*t*) = *θ*(*t*) − *θ* gives 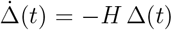. Component-wise, 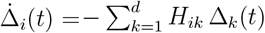. If *H*_*ij*_≠ 0 for some *j* ≠ *i*, then 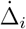 depends on Δ_*j*_, hence the coordinates are coupled.

[OU stationary covariance ⇔ Lyapunov equation] Consider the linear SDE (Ornstein–Uhlenbeck) in parameter space

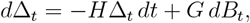

with *H* ∈ ℝ^*d×d*^ Hurwitz (spec(*H*) in the open right half-plane) and diffusion *Q* := *GG*^⊤^ ⪰ 0 constant. If a stationary Gaussian law exists with covariance 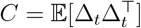, then

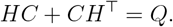

Conversely, if *H* is Hurwitz and *Q* ⪰ 0, there is a unique *C* ⪰ 0 solving the Lyapunov equation, and the OU process is stationary Gaussian with covariance *C*.

Apply Itô to 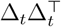:

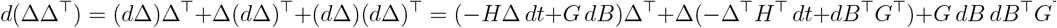

Taking expectations and using 𝔼[*dB*] = 0, 𝔼[*dB dB*^⊤^] = *I dt*:

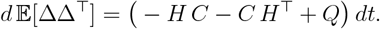

At stationarity *d* 𝔼[ΔΔ^⊤^] = 0, giving *HC* + *CH*^⊤^ = *Q*. Conversely, if *H* is Hurwitz, the continuous Lyapunov equation has a unique *C* ⪰ 0, and the OU process admits the stationary Gaussian with that covariance (standard OU theory).

[Weight–representation correspondence under local Boltzmann/OU regime] Let *a*_*ℓ*_(*θ*_*ℓ*_, *x*) be the activation vector at layer *ℓ* for input *x*, and fix a distribution over inputs with mean-zero feature 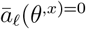 (or subtract the empirical mean). Assume within an epoch the parameter fluctuation Δ_*ℓ*_ := *θ*_*ℓ*_ −*θ*_*ℓ*_ follows the OU approximation with stationary covariance *C*_*ℓ*_ (Lemma 12). If *a*_*ℓ*_ is differentiable in *θ*_*ℓ*_ and we linearize in a neighborhood of *θ*_*ℓ*_,

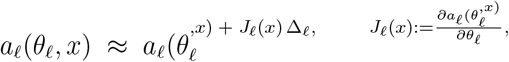

then the representational covariance (over both parameter fluctuations and inputs) satisfies

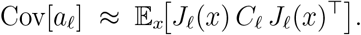

In particular, if *J*_*ℓ*_(*x*) varies weakly with *x* (or we replace it by *J*_*ℓ*_ := 𝔼_*x*_[*J*_*ℓ*_(*x*)]), then 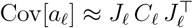.

Let 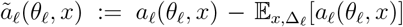. Under the linearization *ã*_*ℓ*_ ≈ *J*_*ℓ*_(*x*) Δ_*ℓ*_ + (zero-mean residual). Taking covariance over the joint law (independent *x* and Δ_*ℓ*_),

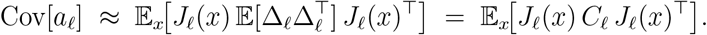

If *J*_*ℓ*_(*x*) is replaced by its mean *J*_*ℓ*_ (e.g., by linear response or a fixed probe set), the approximation reduces to 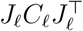.

#### Consequence (entropy–modularity link in representations)

From Theorem 12 and the Gaussian OU covariance *C*_*ℓ*_ (Lemma 12), the activation covariance inherits block structure from *C*_*ℓ*_ through *J*_*ℓ*_. Under fixed marginal variances, increasing within-block correlations (higher modularity) decreases det Cov[*a*_*ℓ*_] by Hadamard’s inequality, thus lowering differential entropy of *a*_*ℓ*_. This matches the parameter-space result and provides the observable bridge.

## Glossary

ViT: Vision Transformer. A transformer-based architecture for image analysis composed of patch embeddings, multi-head attention, and feed-forward layers.
WSI: Whole Slide Image. Gigapixel histopathology slides digitized for computational analysis.
TCGA: The Cancer Genome Atlas. A large cancer genomics consortium providing genomic, transcriptomic, epigenetic, and imaging data.
BRCA: TCGA Breast Invasive Carcinoma cohort. A specific TCGA dataset containing breast cancer molecular and imaging data.
SGD: Stochastic Gradient Descent. An optimization method using mini-batch estimates of the gradient.
OU Process: Ornstein–Uhlenbeck Process. A stochastic process that models mean-reverting diffusion; used to approximate local weight dynamics.
SDE: Stochastic Differential Equation. A differential equation incorporating both deterministic and random noise terms to model parameter evolution.
FP Equation: Fokker–Planck Equation. A partial differential equation describing the time evolution of the probability density of a stochastic process.
CCA: Canonical Correlation Analysis. A method for comparing representations by finding correlated linear combinations of variables.
SVCCA: Singular Vector Canonical Correlation Analysis. A version of CCA combining singular-value decomposition and CCA to compare neural representations.
PWCCA / PKCCA: Projection-Weighted Canonical Correlation Analysis. A representation-similarity method that weights canonical correlations by variance contribution to estimate drift.
CKA: Centered Kernel Alignment. A similarity metric between layer representations, commonly used to detect representational changes.
OU: Ornstein-Uhlenbeck (listed above but included for clarity). Used for modeling weight diffusion under local linearization.
SNR: Signal-to-Noise Ratio (if appearing anywhere in your spectral sections). Measure of spectral or activation clarity relative to noise.
FFT: Fast Fourier Transform. Algorithm for computing Fourier transforms used in spectral analysis of filters.
ID: Identity Drift. Term used to describe neuron-level representational drift across epochs.
CI: Confidence Interval. Used in bootstrap analysis to quantify variability of network statistics.
RT: Replication Timing. A genomic measure of the temporal order in which DNA replicates during the cell cycle; predicted from WSI.

## References

1. Bengio, Yoshua, et al. Representation Learning: A Review and New Perspectives. 1206.5538 [cs], 2014.

2. Toosi, Tahereh. Representational Constraints Underlying Similarity between Task-Optimized Neural Systems. 2312.08545 [cs], 2023.

3. Mandt, Stephan, Matthew D. Hoffman, and David M. Blei. A Variational Analysis of Stochastic Gradient Algorithms. Proceedings of the 33rd International Conference on Machine Learning, 2016.

4. Welling, Max, and Yee Whye Teh. Bayesian Learning via Stochastic Gradient Langevin Dynamics. Proceedings of the 28th International Conference on Machine Learning, 2011.

5. Ba, Jimmy, et al. Do Deep Nets Really Need to Be Deep? Advances in Neural Information Processing Systems, 2014.

6. Dosovitskiy, Alexey, et al. An Image Is Worth 16×16 Words: Transformers for Image Recognition at Scale. International Conference on Learning Representations, 2021.

7. Raghu, Maithra, et al. SVCCA: Singular Vector Canonical Correlation Analysis for Deep Learning Dynamics and Interpretability. Advances in Neural Information Processing Systems, 2017.

8. Maslov, Sergei, and Kim Sneppen. Specificity and Stability in Topology of Protein Networks. Science, vol. 296, no. 5569, 2002, pp. 910–913.

9. Newman, Mark E. J., and Michelle Girvan. Finding and Evaluating Community Structure in Networks. Physical Review E, vol. 69, no. 2, 2004, p. 026113.

10. Güçlü, Umut, and Marcel A. J. van Gerven. Deep Neural Networks Reveal a Gradient in the Complexity of Neural Representations across the Ventral Stream. Journal of Neuroscience, vol. 35, no. 27, 2015, pp. 10005–10014.

11. Morcos, Ari S., et al. On the Importance of Single Directions for Generalization. International Conference on Learning Representations, 2018.

12. Mandt, Stephan, Matthew D. Hoffman, and David M. Blei. A Variational Analysis of Stochastic Gradient Algorithms. Proceedings of the 33rd International Conference on Machine Learning, 2016.

13. Welling, Max, and Yee Whye Teh. Bayesian Learning via Stochastic Gradient Langevin Dynamics. Proceedings of the 28th International Conference on Machine Learning, 2011.

14. Raghu, Maithra, et al. SVCCA: Singular Vector Canonical Correlation Analysis for Deep Learning Dynamics and Interpretability. Advances in Neural Information Processing Systems, 2017.

15. Kornblith, Simon, et al. Similarity of Neural Network Representations Revisited. Proceedings of the 36th International Conference on Machine Learning, 2019.

16. Blondel, Vincent D., et al. Fast Unfolding of Communities in Large Networks. Journal of Statistical Mechanics: Theory and Experiment, 2008, P10008.

17. Newman, Mark E. J., and Michelle Girvan. Finding and Evaluating Community Structure in Networks. Physical Review E, vol. 69, no. 2, 2004, p. 026113.

18. Efron, Bradley, and Robert J. Tibshirani. An Introduction to the Bootstrap. Chapman & Hall, 1993.

19. Kornblith, Simon, et al. Similarity of Neural Network Representations Revisited. Proceedings of the 36th International Conference on Machine Learning, 2019.

20. Chung, Fan R. K. Spectral Graph Theory. American Mathematical Society, 1997.

21. Kuhn, Harold W. The Hungarian Method for the Assignment Problem. Naval Research Logistics Quarterly, vol. 2, no. 1–2, 1955, pp. 83–97.

22. Dietzen, Michelle, et al. Replication Timing Alterations Are Associated with Mutation Acquisition during Breast and Lung Cancer Evolution. Nature Communications, vol. 15, no. 1, 2024.

23. Dietzen, Michelle, et al. Replication Timing Alterations Are Associated with Mutation Acquisition during Breast and Lung Cancer Evolution. Nature Communications, vol. 15, no. 1, 2024.

24. Dietzen, Michelle, et al. Replication Timing Alterations Are Associated with Mutation Acquisition during Breast and Lung Cancer Evolution. Nature Communications, vol. 15, no. 1, 2024.

25. Raghu, Maithra, et al. SVCCA: Singular Vector Canonical Correlation Analysis for Deep Learning Dynamics and Interpretability. Advances in Neural Information Processing Systems, 2017.

26. Fortunato, Santo. Community Detection in Graphs. Physics Reports, vol. 486, no. 3–5, 2010, pp. 75–174.

27. Kornblith, Simon, et al. Similarity of Neural Network Representations Revisited. Proceedings of the 36th International Conference on Machine Learning, 2019.

28. Pavliotis, Grigorios A. Stochastic Processes and Applications: Diffusion Processes, the Fokker–Planck and Langevin Equations. Springer, 2014.

29. Bracewell, Ronald N. The Fourier Transform and Its Applications. McGraw-Hill, 2000.

30. Shannon, Claude E. A Mathematical Theory of Communication. Bell System Technical Journal, vol. 27, 1948, pp. 379–423, 623–656.

